# Joint attention biases dogs’ memory towards object identity

**DOI:** 10.1101/2025.11.14.687955

**Authors:** Andrea Sommese, Maleen Thiele, Christoph J. Völter

## Abstract

Ostensive communication, characterized by direct eye contact and other attention-catching signals, shapes how humans encode and remember novel objects, biasing memory toward identity-relevant features. Dogs are sensitive to human ostensive cues and follow gaze direction. However, whether these signals also enhance and modulate object encoding remains unknown. In an eye-tracking study, we tested dogs in a violation-of-expectation paradigm. A human actor directed eye contact toward the dog or looked away while looking at an object. After an occlusion, dogs viewed three outcomes: the same object (no change), the same object in a different location (location change), or a different object (identity change). We measured dogs’ looking times during the outcome phase within predefined areas of interest around the object. In line with our predictions and matching earlier findings with infants, dogs looked longer at identity changes in the eye-contact condition. In contrast, looking times at location and no-change outcomes were unaffected by communicative context, indicating selective enhancement of identity encoding. During the initial addressing phase, pupil dilation was greater in the eye-contact condition, indicating increased arousal or engagement. These findings demonstrate that dogs and infants exhibit a communication-induced memory bias, revealing a capacity for ostension-guided learning that facilitates information transfer.

## Introduction

Human infants possess a remarkable capacity to acquire culturally shared knowledge through social learning mechanisms that emerge within their first year of life [1,2]. Among the key mechanisms is joint attention, the shared and mutually recognized attention between two individuals and an object [1–5]. Joint attention is often established through ostensive signals, such as eye contact, verbal addressing [6,7] or physical proximity and touch [8–11]. The role of joint attention extends beyond attentional coordination to enhancing memory formation and learning. From four months onwards, infants show enhanced object encoding when they observe an adult looking at an object after being addressed ostensively [12–14], with their sensitivity to eye contact increasing across infancy (live interactions: 14, 15; screen-based settings: 16–18).

As infants progress into the second half of their first year, the influence of joint attention and communicative context on their memory formation becomes even more evident: they show selective encoding of identity-relevant object features [19–21]. This “communication-induced memory bias” aligns with predictions derived from the Natural Pedagogy framework, which assumes that communicative signals bias the learner toward object features that are relevant for the learning of kind-relevant, generalizable information [22–24]. Initial evidence for this bias in preverbal infants came from a study by Yoon et al. [21]. In a screen-based setup, 9-month-olds who encountered an object in a joint attention interaction involving communicative cueing by an adult (eye contact, infant-directed speech, pointing), later looked longer when the object’s identity had changed than when its location had changed, suggesting selective encoding of internal, identity-relevant over external, transient object features. Although not replicated in one study [25] subsequent work has demonstrated similar patterns using only eye contact to establish the communicative context in live interactions [20] and in screen-based settings [19]. The latter study also provided evidence that the bias can arise through third-party observation [19]. A similar communication-induced memory bias has also been found in human adults [26,27].

Despite extensive research on human social learning, the evolutionary origins and cross-species generality of ostension-guided learning remain largely unexplored [28]. Dogs represent an ideal comparative model, given their unique evolutionary history with humans and their demonstrated sensitivity to human communicative signals [29–39]. Canine cognition research has shown that dogs track human attentional states [40,41], exhibit visual perspective-taking abilities [29–31,35,42] and even monitor human knowledge states [33,34,38]. Ostensive cues such as verbal addressing and eye contact reliably enhance dogs’ gaze following and bias their choices [32,33,36,37,39,43,44]. Importantly, dogs also form durable object representations in social contexts [e.g., 45,46] and can track object identity across occlusion and delay [47,48], raising the possibility that communicative cues may influence not only immediate orienting but also what information is encoded into memory. What remains less clear is whether these cues shape not only where dogs attend but also what they encode about objects, specifically, whether eye contact induces a human-like bias to prioritize identity over location. To our knowledge, only one study has recently investigated the influence of communicative context on canine object memory [49]. Using a live-interaction paradigm, the authors found that dogs generally looked longer at identity changes than location or no-change controls, regardless of the communicative context. However, because eye contact, ostensive speech, and pointing were collapsed into a single communicative condition (replicating the design of Yoon et al., 2008 [21]), the specific contribution of eye contact—independent of other cues—remains unknown.

Here, we tested this question in a preregistered eye-tracking study using the violation-of-expectation paradigm and video stimuli by Thiele et al. [19]. Pet dogs viewed videos in which a woman either looked toward the dog (eye-contact condition) or looked away (no-eye-contact condition) before and after looking at an object. Following a brief occlusion, we presented one of three outcomes: the same object reappeared in the same location on screen (no change), the same object appeared in a different location (location change), or a completely novel object appeared in the same location (identity change). If dogs share humans’ preparedness for ostension-guided learning, we predicted that they should selectively encode recognition-relevant object properties and, consequently, look longer to identity changes, particularly following eye contact.

## Material and methods

The experimental design, hypotheses, predictions, sample size, and analysis plan were preregistered (https://doi.org/10.17605/OSF.IO/YZ6AP). All analysis scripts and data associated with the study are openly available in the repository cited in the Data, Materials, and Software Availability section. The procedure described in this study was discussed and approved by the Ethics and Animal Welfare Committee of the University of Veterinary Medicine Vienna, following the university’s guidelines for good scientific practice (ETK-179/11/2023).

### Subjects

Our target sample size (N = 36) was determined based on the results of an a priori power simulation reported by Thiele et al. [19], which used the same design and stimuli in a study with human infants. The simulation estimated effect sizes using raw data from Okumura et al. [20]. We started testing 36 pet dogs (18 females) of various breeds and mixes (Table S5 provides demographic information). The mean age at the beginning of the experiment was 56.1 months (±4.21; range 11–103 months). Of these, 34 dogs completed all trials, while one additional dog became unavailable after the first session but was still included in the analyses (final sample: N = 35). Dogs had no prior experience with the specific stimuli or task used in this study.

### Study design

Following the pretest, each dog completed 12 test trials structured according to a 2 × 3 factorial design (using the design by 18). The two manipulated factors were: (1) the presence of eye contact between the actor and the dog during the action phase (eye contact vs. no eye contact) and (2) object change during the outcome phase (no change, identity change, location change). Each dog experienced two trials per condition, for a total of six conditions. Trials were distributed across four sessions, each consisting of three trials with distinct outcome conditions.

### Stimuli

We used the video stimuli created for the study by Thiele et al. (18), to allow direct comparison with infant data, differing only in our selection of object images, which were taken from the same object collection that was initially created for a study by Wahl et al. [14]. Each video consisted of 3 phases: (1) an action phase featuring a 15-second video of an actor gazing at a novel object; (2) a delay phase in which the scene was occluded for 3 seconds; and (3) an outcome phase where a single object was shown for 15 seconds or until the dog looked away from the screen for two consecutive seconds, following a minimum of 100 ms of gaze at the object’s area of interest (AoI; 350 × 350 px).

During the **action phase**, a curtain opened (1s) to reveal one of two adult women on either side of the screen (actor AoI: 550 × 760 px). The actor’s position (left or right) was fully counterbalanced across dogs, eye-contact conditions, and outcome conditions. Each dog saw the actor three times on each side within each condition. For the first two test trials, the actor’s position was counterbalanced so that half of the dogs saw her on the left first, and the other half on the right, with the opposite position in the second trial. The actor was initially shown in back view (1s). In the eye-contact condition, the actor first turned toward the dog (1s) and looked at the dog for (1s). Then, she shifted her gaze during 1s to a centrally visible object (positioned at the top or bottom of the screen, counterbalanced), looked at it (1s), and returned her gaze to the dog during 1s, looking again at the dog (1s). This alternating gaze shift between the dog and the object was repeated twice. In the no-eye-contact condition, timing and body movements were identical, but the actor was seen in a side-view, facing outward and never establishing eye contact with the dog. Instead, she alternated her gaze between the object and her original side-facing position. In both conditions, the actor maintained a neutral facial expression and did not speak. The action phase ended with the curtain closing (1s). During the delay phase, the screen remained occluded by the curtain (3s). Finally, in the outcome phase, the curtain reopened (1s) to reveal one of three outcomes: (1) the same object in the same screen location (no change), (2) the same object in a new location (location change), or (3) a different object in the same location (identity change). For a detailed illustration of the timing of the experimental phases, see Figure 1 in Thiele et al. [19]. For an example video illustrating the temporal sequence of a single trial, see Movie S1 in the Supplementary Materials.

**Figure 1.**
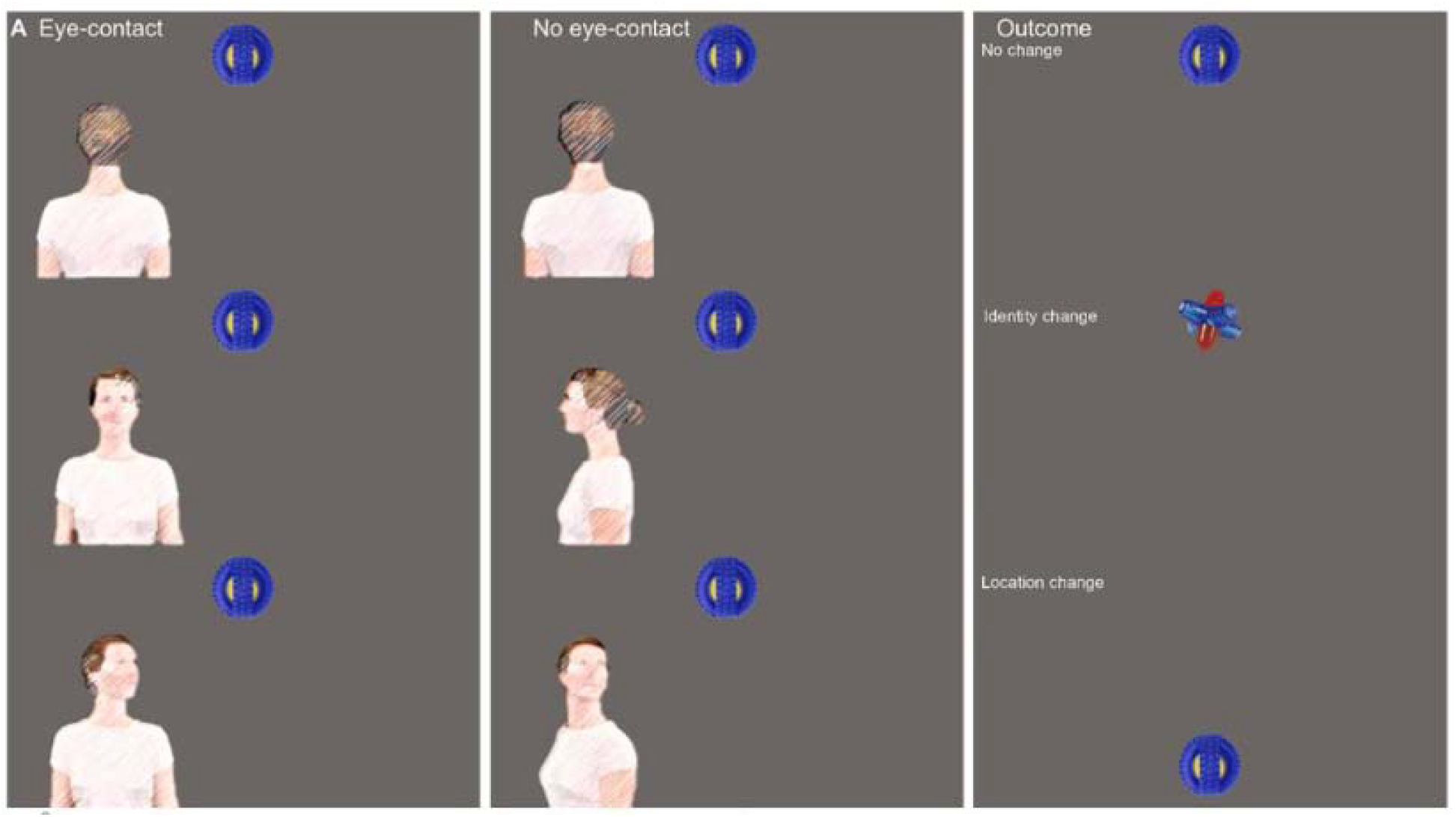
Screenshots of the video stimuli in the eye-contact (left) and no-eye-contact (center) conditions of the still phases in the action phase and of the outcome conditions in the outcome phase (right). The positioning of the actor (left or right on screen) and of the object in the action phase (top or bottom) wa counterbalanced. *Based on stimuli originally developed for Thiele, M*., *Kalinke, S*., *Michel, C*., *& Haun, D. B. (2023). Direct and observed joint attention modulates 9-month-old infants’ object encoding. Open Mind, 7, 917–946*.

**Figure 1.**
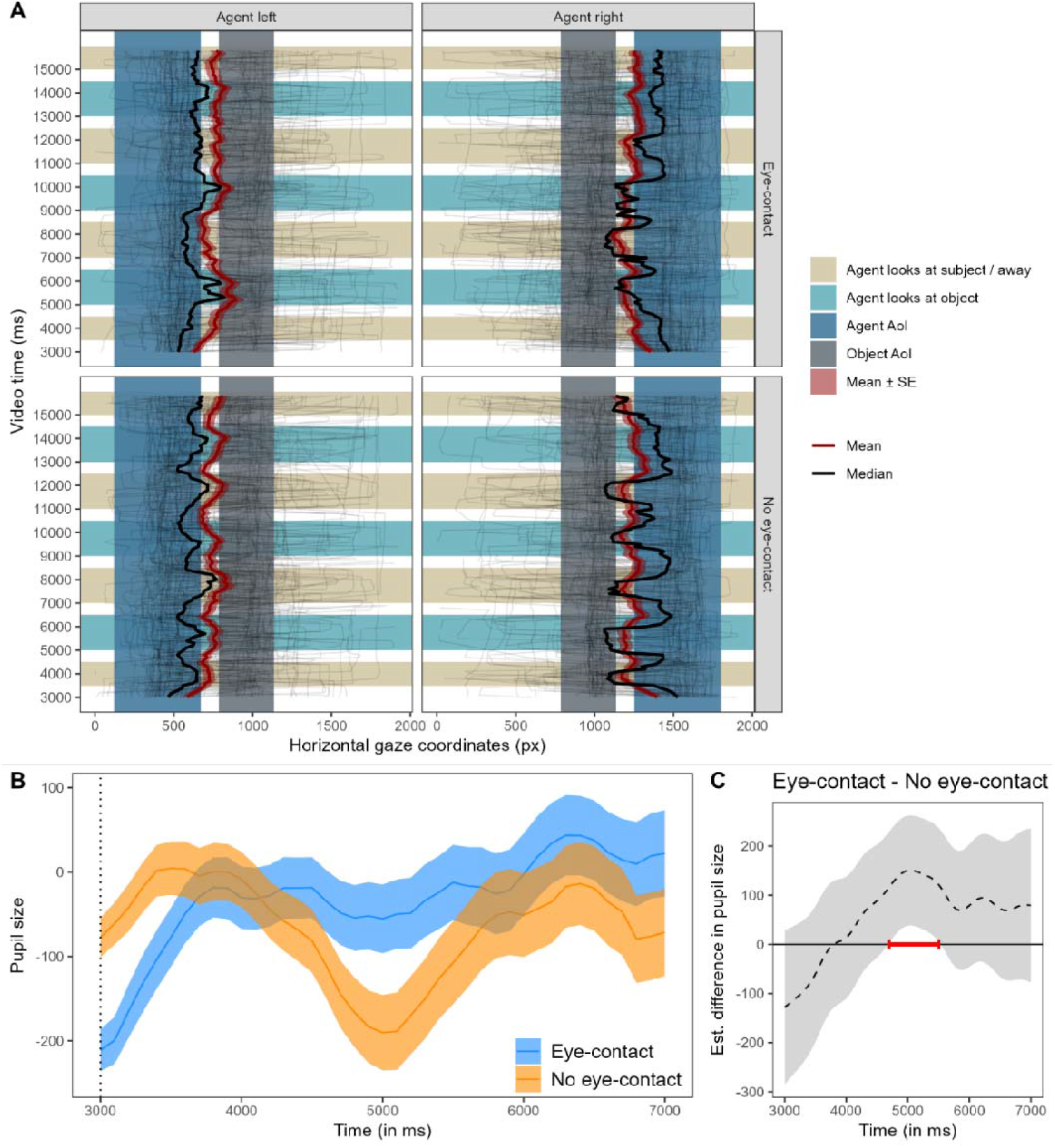
Action phase gaze and pupil size visualizations. (A) Gaze time course during the action phase. Shown are dogs’ horizontal gaze coordinates (px) over time for the eye-contact and no-eye-contact conditions, split by the actor’s screen position (left vs right). Thin lines depict individual gaze trajectories; thick lines depict the grand mean (dark red) and median (black), with shaded bands indicating ±1 SE around the mean. Background shading marks predefined temporal segments in which the actor looked toward the subject (eye-contact condition) / away (no eye-contact condition; orange) or looked at the object (green). Additional shaded bands indicate the predefined agent and object areas of interest (AoIs) along the X axis. (B) Time series plot showing dogs’ pupil size (in arbitrary units and baseline corrected). The blue and orange lines show the mean pupil size (± SE, shaded area around the line) in the action phase of the eye-contact and no-eye-contact conditions. The dashed vertical line (at 3000 ms) indicate when the actor turns toward the camera in the eye-contact condition. (C) Difference curve obtained from GAMM01. The dashed line shows the estimated difference between the eye-contact and no-eye-contact conditions during the action phase, and the shaded area shows the pointwise 95% confidence interval. The period in which the conditions differ significantly is highlighted in red.

During the first testing session, dogs participated in two pretest trials before the main experiment started. These trials were designed to familiarize the dogs with the depicted actors, the structured sequence of action, occlusion, and outcome phases used in the test trials, and the two possible object positions [19–21,25]. The pretest videos followed the same temporal structure but without referential gaze shifts during the action phase. Instead, the actors turned either toward or away from the dog. The outcome phase always showed the same object in the same location. The actor in the eye-contact pretest was the same as in the eye-contact test trials, and likewise for the no-eye-contact condition.

Each dog saw two different actors, one for each condition, and the assignment of actors to conditions was counterbalanced across subjects. Sixteen toy images (260×260 px) were used during test trials, counterbalanced across the six experimental conditions (2 eye contact × 3 outcome types), such that each object served both as a familiar and a novel item across participants. Two additional objects were used in the pretest phase. The toys were selected from a validated set used in prior infant studies [14,19], ensuring comparability with human data. Additionally, we matched the luminance of the object images by scaling the RGB values of non-transparent pixels to a common target value. We selected visually distinct toys that included common features found in typical dog toys (e.g., rope texture, plush material, rubber shapes). Object screen positions, either top or bottom, were also counterbalanced within and across dogs and conditions to ensure balanced exposure. In the first session, outcome types and social conditions were fully counterbalanced across dogs. In subsequent sessions, trial orders were pseudo-randomized with constraints to prevent more than two consecutive trials of the same type and to ensure all three outcomes appeared once per session.

### Procedure

We used an EyeLink 1000 eye-tracking system (SR Research, Canada) to examine dogs’ gaze behavior in response to the video stimuli. The dogs were trained to place their heads on a chinrest before participating in the experiment; details about the training are provided in Karl et al. [50]. During the training and before each test run, each dog performed 5-point calibrations with animated, non-social targets (size: 64 × 64 px; e.g., a moving fly or cartoon figure). The training was considered successful if dogs achieved a calibration validation with an average of less than 1° of visual angle deviation and maintained their head on the chinrest for at least one minute when the trainer left the room. During training and testing, each dog rested its head on an adjustable chinrest tailored to their body size to ensure a stable head position and consistent viewing distance (approximately 67 cm) from the monitor. Videos played at 25 fps on a 1920□×□1080 px LCD monitor, covering 43.4° by 25.2° of visual angle. Interested owners were allowed to remain in the room during the experiments and watch the screen from behind their dog’s back, approximately two meters from the dog.

At the beginning of each trial, an attention-grabbing animation (identical to the one used during the calibration and different from the one used in the human infant study by Thiele et al. [19]) was presented at the center of the screen to direct and center the dogs’ gaze. The animation preceding the action phase was gaze-contingent, meaning that the trial would only begin if the dog’s gaze were detected for 50 msec within a predefined area of interest (AoI) surrounding the animation (concentric AoI, diameter = 300 px). This procedure ensured engagement and reduced data loss due to inattention. After each video presentation, a gray screen appeared until the following fixation animation or the end of the experiment. If a trial needed to be terminated before the video concluded, such as when the dogs left the chinrest, that trial, and any subsequent trials, were repeated after recalibrating, either in the same session or in a subsequent one (this happened in six trials). During the whole session, water was available to the dogs.

### Statistical analysis

We fitted Generalized Linear Mixed Models (GLMM) with gamma error structure and Linear Mixed Models (LMM) to analyze the dogs’ looking times into the object AoI (AoI size: w x h: 350 × 350 px), latency to look at the object AOI, and first look duration. Following our preregistered analysis plan, we fitted gamma-distributed GLMMs unless the model assumptions were violated. We checked whether the model assumptions were violated using the ‘plotQQunif’ (QQ-plot of residuals) and ‘plotResiduals’ (residuals plotted against predicted values) functions of the DHARMa package [51]. Significant violations concerning the distribution of the residuals were detected for the looking time model, but not for the first look duration model. Therefore, we fitted an LMM for the overall dwell time data instead (and evaluated the Gamma model in the case of the first look duration). We analyzed the overall dwell times as within-subject z-scores to account for individual differences in absolute looking times and to ensure that the assumptions of the LMM were met (e.g., 57). Following the z-transformation, we found no apparent deviations from model assumptions concerning normal distribution and homoscedasticity of residuals. Our primary analyses were based on fixations as determined by the default Eyelink event parsing algorithm, but we also repeated our confirmatory analyses based on the raw samples (excluding blink artefacts; see ESM).

As predictor variables, we include the eye-contact condition (eye-contact or no-eye-contact), outcome type (no-change, identity-change, location-change) and their interaction, as well as the object position in the outcome phase (top or bottom) and overall trial number (z-transformed). The session number and the number of trials per session varied across dogs. Therefore, we included the overall trial number instead of the session number and trials within the session. Dog ID was included as a random effect as well as (centered) random slope components of eye-contact condition, outcome, object position, and trial number to avoid pseudo-replication and to keep the type-I error rate at the nominal level of 5% [53]. In the case of convergence issues, we removed the correlation between random intercepts and random slopes. We evaluated the significance of the interaction by using a likelihood ratio test. Following a significant interaction, we tested the difference between the identity-change vs no-change and location-change vs no-change for each condition of individual fixed effects employing the Satterthwaite approximation using the function lmer of the package lmerTest and a model fitted with restricted maximum likelihood.

For the exploratory analysis, including latency to first fixation within the object AoI (outcome phase), looking time to the object (action phase), and looking time to the actor (action phase; AoI size: w x h: 550 × 720 px), we fitted LMMs based on within-subject z-scores. The model structure was the same as in the confirmatory models (except that the outcome condition was, of course, not added for the action phase models).

All statistical analyses were conducted in R (59; version 4.4.2) using the packages glmmTMB (version 1.1.10; 59), lme4 (version 1.1-35.5; 60), and lmerTest [57]. We obtained confidence intervals for model estimates and fitted values using a parametric bootstrap (N = 1,000 bootstraps). We assessed potential collinearity using Variance Inflation Factors (VIF; 62), which were calculated for a linear mixed model without the interaction term and random effects. The results indicated that collinearity was not a concern, with the maximum VIF being <1.1.

Thirty-six dogs started the experiment, but only 34 completed all test sessions. One additional dog completed the first test session (included); another dog began with the pretest but became unavailable before the start of the test phase (not included). Out of the 409 recorded test trials, we excluded trials if the dogs’ gaze was detected in less than 70% of the action phase video (N=8) and if subjects looked at the object for less than 100 msec in the outcome phase (N=115). Given the dogs’ bias to look at the upper half of the screen and the fact that there was no gaze-contingent onset of the outcome phase, we also filtered trials in which the dogs were looking at the object’s outcome location at the start of the outcome phase and, therefore, were not actively shifting their gaze to the object once it appeared (N = 39). The final sample consisted of a total of 247 trials of 35 dogs.

### Data, Materials, and Software Availability

All data, stimuli, and analysis scripts used in this study are publicly available on Zenodo (https://zenodo.org/records/17553617). The repository contains raw and pre-processed eye-tracking data, stimulus materials and all R scripts required to replicate the pre-processing, modelling and statistical analyses described in the manuscript. There are no restrictions on data access.

## Results

To examine whether ostensive signals influence dogs’ object encoding, we analyzed dogs’ looking patterns and pupil size changes in an eye-tracking experiment using a violation-of-expectation paradigm (Fig. 1). Confirmatory and exploratory analyses revealed that dogs looked longer and faster at objects that changed identity, specifically following eye contact. These effects were not observed for location changes or unchanged objects. Additional analyses showed increased pupil dilation during eye contact, supporting heightened arousal or engagement.

### Action phase

#### Dwell times to actor and object in the action phase (exploratory analysis)

During the action phase, dogs looked back and forth between the agent and the cued object in both conditions (Fig. 2A). To ensure that our results reflected differential encoding rather than biased attention during initial stimulus presentation, we analyzed the dogs’ looking behavior during the action phase when the human actor looked at the object. We found no evidence for a significant difference in attention during the action phase between conditions, neither in the dogs’ looking times to the actor (χ2(1) = 0.54, p = 0.461) nor the object (χ2(1) = 0.65, p = 0.422).

**Figure 2.**
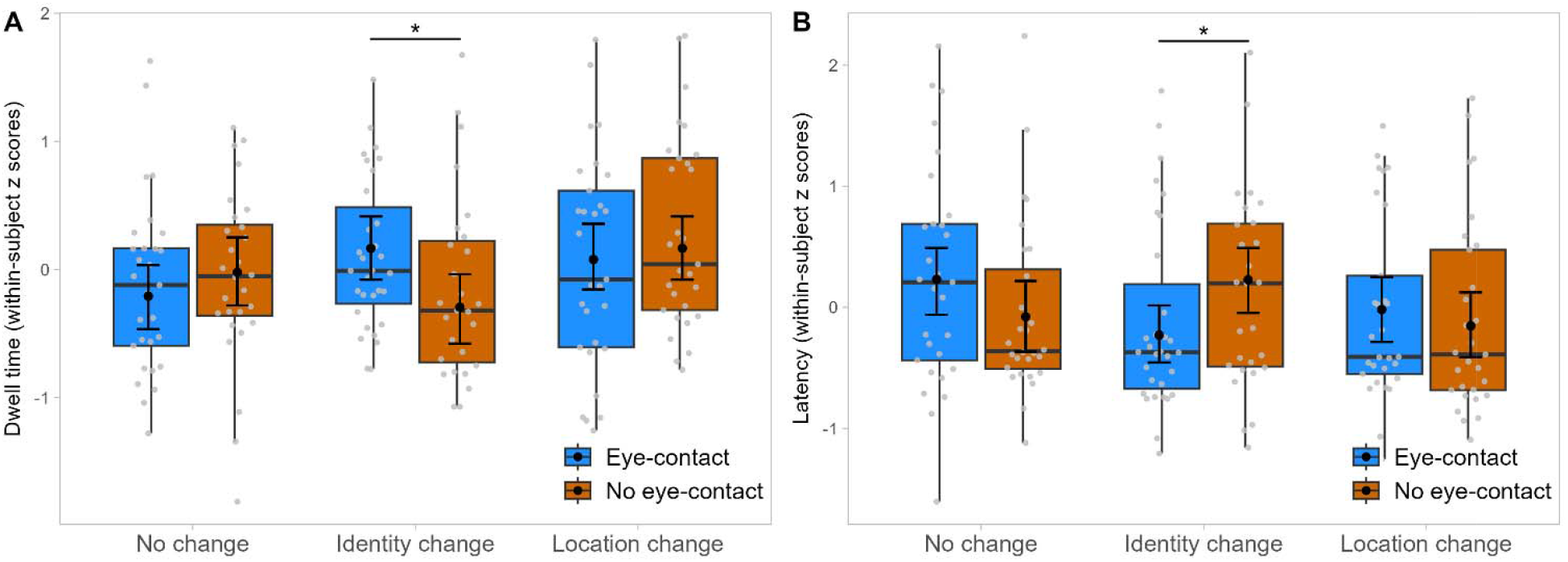
Outcome phase area of interest analysis. (A) Boxplot showing the dogs’ dwell time (within-subject z-scores) in the object AoI during the outcome phase across the eye-contact and outcome conditions. (B) Latency (shown as within-subject z-scores) of the first look into the object AoI in the outcome phase. Data are grouped by condition, with eye-contact trials in blue and no-eye-contact trials in orange. Grey dots represent the mean values of each subject, while black dots show the fitted model, and the error bars are its 95% confidence interval: *p < 0.05.

#### Pupil size changes in the action phase (exploratory analysis)

We analyzed the preprocessed, baseline-corrected pupil size data at the beginning of the action phase, in which the actor turned around looking toward the dog’s direction or to the side (Fig. 2B; for the raw pupil size data, see Fig. S1 A-B) using a generalized additive mixed model (GAMM01). The full model, including condition, fitted the data significantly better than the null model (Chi-square test of ML scores: χ2(3) = 9.51, p < 0.001; GAMM01 had a lower AIC: ΔAIC 26.96). The pupil size varied significantly over time in both conditions (eye-contact: F(12.67, 14.88) = 4.74, p < 0.001; no eye-contact: F(13.14, 15.32) = 3.42, p < 0.001). While over the entire interest period (3000 – 7000 ms) there was no significant difference between the dogs’ pupil size in the eye-contact and no eye-contact condition (t = - 1.10, p = 0.271), the difference curve revealed that the pupil size was significantly larger in the eye-contact condition than the no-eye-contact condition in the time window between 4697 and 5505 ms (Fig. 2C; for the model estimates and partial and summed effects see Table S4 and Fig. S2).

### Outcome phase

#### Dwell time in the outcome phase (confirmatory analysis)

Dogs spent significantly more time looking at objects when their identity changed, but only after the human actor had made eye contact with them (Fig. 3A). The full model provided a significantly better fit to the data compared to the null model, which only included the random effects (χ2(7) = 30.80, p < 0.001). As predicted, we found a significant interaction between condition and outcome (χ2(2) = 7.08, p = 0.029; Table S1). The difference between the eye-contact and no-eye-contact conditions in the dwell times was significantly greater in the identity change than in the no-change condition (t = -2.19, df = 217.53, p = 0.029). No significant interaction was found for the location change vs no change contrast (t = -0.11, df = 213.88, p = 0.909). We found that the dogs looked significantly longer at the object in the eye-contact than no-eye-contact condition following an identity change (t = -2.35, df = 215.19, p = 0.020) but not following a location change (t = 0.64, df = 211.93, p = 0.524) or no change (t = 0.76, df = 212.04, p = 0.448; see ESM for mean dwell times). Additionally, the dogs looked significantly more often at the upper object (t = 4.81, df = 33.88, p < 0.001). The trial number had no significant effect (t = 0.85, df = 30.96, p = 0.400).

#### Latency to look at the object in the outcome phase (exploratory analysis)

Dogs not only looked longer at identity changes following eye contact but also oriented to these changes more rapidly (Fig. 3B). The full model provided a significantly better fit to the data compared to the null model, which only included the random effects (χ2(7) = 22.54, p < 0.001). As before, we found a significant interaction between condition and outcome (χ2(2) = 8.87, p = 0.012; Table S2). The difference between the eye-contact and no-eye-contact conditions was significantly greater in the identity change than in the no-change condition (t = -2.49, df = 179.44, p = 0.014). No significant interaction was found for the location change vs no change contrast (t = -0.53, df = 181.31, p = 0.600). We found that the dog looked significantly faster at the object in the eye-contact than no-eye-contact condition following an identity change (t = 2.10, df = 127.65, p= 0.038) but not following a location change (t = -0.63, df = 122.47 p = 0.533) or no change (t = -1.31, df = 134.40, p = 0.194). Additionally, the dogs looked significantly sooner at the upper object (t = -3.95, df = 27.51, p < 0.001). The trial number had no significant effect (t = -1.76, df = 208.10, p = 0.080).

## Discussion

Our findings provide initial evidence that ostensive signals, specifically eye contact, selectively influence dogs’ encoding of object identity, mirroring communicative learning biases reported in human infants [19–21]. The pet dogs in our study oriented more rapidly and looked longer at identity changes, but only when the actor first established eye contact. This suggests that, as in infants, ostensive cues direct information processing toward intrinsic features of objects rather than transient spatial information.

These findings show that dogs’ memory and attention can be shaped not merely by associative processes or stimulus salience, but also by communicative context. This effect is closely aligned with observations in preverbal infants, who also selectively encode identity-relevant properties of objects when interactions are marked by ostensive signals and interpersonal jointness [19–21]. Within this broader framework, our results are consistent with predictions derived from the Natural Pedagogy theory [22,23] beyond humans and hominids [28]. The theory originally proposed that humans are specifically prepared to use ostensive signals from an early age to learn generalizable information from others. Our data suggest that at least one domesticated nonhuman species exhibits similar selectivity, though this does not necessarily mean they share the same learning mechanisms. These findings are consistent with the notion that domestication and enculturation with humans have tuned dogs to treat direct eye contact in combination with head turns toward an object as cues to relevant and generalizable content as well.

Unlike the study by Johnston et al. [49] in which dogs looked longer at identity changes in both communicative and non-communicative conditions, our results revealed a selective response to identity changes only following eye contact. This difference may reflect methodological factors: Johnston et al. relied on live interactions and manual coding of looking times, whereas we used tightly controlled video stimuli and automated eye tracking, allowing precise timing, consistent condition matching, and higher temporal and spatial resolution of the looking-time measure, as well as pupillary measures. Additionally, Johnston et al. operationalized communicative context using multiple cues—such as pointing or reaching gestures and name-calling—making it difficult to isolate the specific contribution of eye contact independently. Together, these differences underscore how subtle variations in design and attentional control can substantially influence measured memory biases, as also highlighted in the infant literature [19,25].

In the context of pointing, another study [59] suggested that dogs preferentially encode an object’s location rather than its identity-relevant features following ostensive addressing. To examine this, the experimenter, after pointing at one of the objects, switched the locations of two objects visible to the dogs. After the switch, dogs preferentially approached the initially cued location rather than the originally cued object in the condition involving eye contact and verbal addressing. In contrast, our findings suggest that eye contact can bias dogs’ memory toward recognition-relevant object features. Two non-exclusive explanations could resolve this discrepancy with our findings. First, pointing may be interpreted by dogs as an imperative, spatial directive. If so, pointing might bias dogs toward locations or directions, whereas eye contact alone can bias encoding toward object identity (but see Johnston et al., 2021 [49]). Second, several studies have shown that ostensive communication can induce search errors in dogs even when they have witnessed where an object has actually been hidden [32,33,37,43,44], which could inflate location-based choices. Our paradigm avoided pointing, hiding, and choice demands, and we still observed an identity bias selectively after eye contact. Thus, when isolating ostension and minimizing task demands that potentially emphasize the spatial dimension or response habits, dogs exhibit a human-like bias toward identity-relevant features.

Pupillometry offers a window into potential mechanisms. Eye contact elicited a transient increase in pupil size, consistent with a brief arousal or engagement response, potentially facilitating encoding of referential information into memory. Indeed, in humans, pupil dilation during encoding of new information is predictive of successful later memory retrieval [60,61]. Additionally, in human infants, eye contact can lead to increased arousal (measured using heart rates) and predict subsequent responsiveness to referential cues [62,63].

While these results indicate a parallel between dogs and infants in their sensitivity to ostensive cues, dogs likely still differ from human infants in how flexibly and broadly they generalize information, and the degree to which they rely on familiarity or training [37]. Throughout their domestication, dogs might have undergone selection for socio-cognitive abilities that allow them to interpret and adequately respond to human communicative signals [37,64]. However, more research is needed to elucidate the boundaries and mechanisms underlying this capacity, and how it compares to those of other species, both domesticated and non-domesticated.

Although our study benefits from the accuracy and detailed data offered by eye-tracking technology, some limitations merit consideration. A primary concern is the use of screen-based stimuli, which differs from dogs’ real-life interactions and therefore raise questions about ecological validity. However, evidence from studies with infants shows that communication-induced memory biases emerge both in screen-based procedures [19,21] and in live interactive setups [20], suggesting that the underlying mechanism is consistent across contexts and not an artefact of presentation format. Screen-based research paradigms also indicate that dogs actively engage with and interpret social and communicative signals on screens. For instance, Téglás et al. [36] found that dogs follow human gaze cues presented on a screen, particularly after being addressed ostensively. Karl et al. [65] found that dogs looked longer at pictures of their caregivers on a screen than at another familiar person, and their pupils dilated more when the depicted person exhibited an angry facial expression. Further screen-based research shows that dogs can learn to categorize pictures [66], anticipate the destination of moving objects on the screen [67], react to animacy cues [67], and show sensitivity to certain physical regularities (53, 54; for a review, see 38). Together, these findings support the validity of using screen-based paradigms to investigate cognitive processes in both humans and dogs. In our study, we further controlled for passive viewing by excluding trials where dogs were already looking at the object’s location at the onset of the outcome phase, ensuring that recorded gaze shifts reflected an active response to the social cue rather than passive viewing. Moreover, our experimental design closely followed the paradigm developed by Thiele et al. [19] for infants, employing the same video stimuli and trial structures to enable direct cross-species comparison.

Future research would benefit from testing how dogs’ ostension-guided memory encoding unfolds in more naturalistic settings, using live social partners and incorporating multimodal cues such as gestures, vocalizations, prosody, and body orientation. Understanding how these cues interact, whether they are additive, redundant, or context-dependent, could offer further insight into the mechanisms by which dogs infer relevance and prioritize information. For instance, pointing gestures or vocal intonation may shift dogs’ expectations toward particular types of content (e.g., spatial vs. identity-relevant information), potentially modulating what they encode [59]. Factors such as familiarity, emotional expression, or perceived relevance may affect how ostensive signals are interpreted and how readily they facilitate learning. Comparative studies examining the impact of social versus non-social attention-getters, such as sudden movement or sound, could also help disentangle whether the observed effects are specific to communicative intent or reflect broader attentional processes. Finally, individual differences likely influence how dogs respond to ostensive cues. Factors such as breed, training history, and life experience may shape sensitivity to communicative signals. For instance, working dogs may be more attuned to intentional cues due to training and companion breeds might show heightened responsiveness to eye contact because of selection for social engagement.

## Conclusions

In summary, our results demonstrate that human communicative signals (eye contact) not only capture canine attention but can also bias which features of a scene are encoded and remembered, similar to mechanisms of early social learning seen in humans. These findings have practical implications for developing more effective training programs for pet and working dogs based on communicative engagement and joint attention. They also contribute to theoretical discussions on Natural Pedagogy, suggesting that some mechanisms of ostension-guided learning may be shared at least with a domesticated species. By aligning our experimental design with studies in human infants, we provide a comparative framework to explore the evolutionary origins of social cognition and highlight the potential for dogs to serve as models for understanding cross-species learning mechanisms.

## Acknowledgments

We want to thank the dogs and their caregivers, Laura Laussegger and Marion Umek, for collecting the data, and Karin Bayer for providing administrative support. This research was funded in whole or in part by the Austrian Science Fund (FWF) [grant DOI: 10.55776/P36896]. For open access purposes, the author has applied a CC BY public copyright license to any author-accepted manuscript version arising from this submission.

