## Supplementary material for "Joint attention biases dogs’ memory towards object identity": SI

**Pupil size processing and analysis**

We pre-processed the pupil size data as follows: we removed blink artifacts by extending the detected blinks by 100 ms. We then calculated the speed of pupil diameter changes over time and the median speed values for each recording session. We detected outliers based on the median absolute deviation (MAD) method, with a threshold set at eight times the MAD. Finally, missing values were linearly interpolated (up to gaps of 500 ms). We then conducted a subtractive baseline correction. For the baseline correction, we used the median pupil size in the phase from 2000-3000 ms. We then sampled the data down to 10 Hz (by extracting the median values in 100 ms bins). After the baseline correction and down-sampling, we repeated the MAD-based artefact correction and interpolation steps.

We fitted a generalized additive mixed model (GAMM) with a Gaussian error structure to analyze the pre-processed pupil size data (following the recommendations by 1, 2, and in line with our previous pupil size analyses, see 3, 4). We analyzed a 4-s interest period starting at the end of the baseline period, following the analysis pipeline of our previous pupillometry studies (3).

We implemented the GAMM in R using the function ‘bam’ of the package ‘mgcv’ (5) and package ‘itsadug’ (6) for visualization. We used the smoothing parameter selection method ‘ML’. We included eye-contact condition as a linear term, the nonlinear regression lines for the two levels of condition over time (with the upper limit for the number of basis functions set to 20), and the nonlinear interaction between X and Y gaze positions (given that the gaze position can affect the pupil size) (2, 7). We also included random factor smooths for each subject and for each time series trajectory (i.e. for each subject and test trial) to improve the model fit and to account for autocorrelation (1, 2).

We evaluated the model fit by inspecting visualizations of correlations between the residuals and the lagged residuals, a QQ-plot of residuals, as well as the residuals against the fitted values (using the functions’ gam.check’ of package ‘mgcv’ and ‘acf’ of package ‘stats’). The residuals appeared to be approximately normally distributed, and there was no evident pattern in the residuals plotted against the fitted values.

For evaluating the significance of condition on pupil size, we compared the full model to a reduced model excluding both the parametric and smooth terms of condition using a Chi-square test of ML scores (using the function compareML of the R package ‘itsadug’) (6). Additionally, we inspected the model summary and visually inspected the estimates of the differences between the conditions (using the function plot_diff of the R package ‘itsadug’) (6).

**Looking times (based on the default Eyelink event parsing algorithm)**

*Descriptive statistics*

Dogs looked at the object in the identity change condition, on average, 869 ms (se 717 ms) longer following eye-contact (5721 ms ± 676 ms) than no-eye-contact (4778 ms ± 676 ms). For the other two outcomes, the pattern was reversed. In the location change condition, dogs looked for an average of 958 ms (se 825 ms) longer following no-eye-contact (6069 ms ± 657 ms) than after eye-contact (5465 ms ± 636 ms). Similarly, in the no-change condition, they looked on average 528 ms (SE 678 ms) longer following no-eye-contact (5428 ms ± 582 ms) than after eye-contact (4866 ms ± 592 ms).

**Sample-based looking time analysis**

Analyses with looking times based on samples (not using any event parsing algorithm to quantify fixations). Given that we detected violations of the Generalized Linear Mixed Models (GLMM) with gamma error structure, we again fitted an Linear Mixed Models (LMM) based on within-subject z-scores. For the first look analysis, the assumptions of the gamma model were again met, so we used the gamma model for the first look analysis. We used the same model structure and analysis pipeline as for the dwell time analyses reported in the analysis section.

**Supplementary results using sample-based looking times**

**Confirmatory analysis**

*Looking time (based on samples) in the outcome phase*

The full model provided a significantly better fit to the data compared to the null model, which only included the random effects (*χ*^2^(7) = 27.95, p < 0.001). We found a significant interaction between condition and outcome (*χ*^2^(2) = 6.64, p = 0.036; Table S3). The difference between the eye-contact and no-eye-contact conditions in the dwell times was significantly greater in the identity change than the no-change condition (t = -2.31, p = 0.022). No significant interaction was found for the location change vs no change contrast (t = -0.85, p = 0.397). We found that the dogs looked significantly longer at the object in the eye-contact than no-eye-contact condition following an identity change (t = -2.16, p= 0.033) but not following a location change (t = -0.08, p = 0.933) or no change (t = 1.11, p = 0.269; Fig. 1). Additionally, the dogs looked significantly more often at the upper object (t = 4.63, p < 0.001). The trial number had no significant effect (t = 0.20, p = 0.842).

*First look based on samples*

A comparison between the null model and the full model showed that the full model did not significantly improve the fit compared to the null model (χ²(7) = 8.81, p = 0.266). A reduced model without the interaction term between eye-contact condition and outcome did not significantly improve the model fit either (χ²(5) = 7.25, p = 0.203).

*First object looks in the outcome phase.*

A comparison between the null and full models showed that the full model did not significantly improve the fit compared to the null model (χ2(7) = 12.05, p = 0.099). Furthermore, a reduced model excluding the interaction term between eye-contact condition and outcome did not significantly improve the model fit (χ2(5) = 8.77, p = 0.119).

**Table S1.** Results of the linear mixed-effects model (LMM) analyzing z-scored dwell time on the target object during the outcome phase. The table reports parameter estimates (Estimate ± SE), t values, approximate degrees of freedom (df), and p values from lmerTest, along with 95 % confidence intervals (CIs) and likelihood ratio test (LRT) results.

|  | **Estimate** | **SE** | **t** | **df** | **p** | **LowerCI** | **UpperCI** | **Chi2** | **df_LRT** | **p_LRT** |
| --- | --- | --- | --- | --- | --- | --- | --- | --- | --- | --- |
| **(Intercept)** | -0.59 | 0.14 | -3.64 | 141.2 | <0.001 | -0.85 | -0.31 |  |  |  |
| **Condition^2^** | 0.19 | 0.19 | 0.76 | 212.04 | 0.448 | -0.19 | 0.55 |  |  |  |
| **outcomeIDENTITY** | 0.37 | 0.18 | 1.83 | 218.94 | 0.069 | 0.02 | 0.71 |  |  |  |
| **outcomeLOCATION** | 0.29 | 0.18 | 1.19 | 215.08 | 0.234 | -0.05 | 0.62 |  |  |  |
| **object_pos_outcomeup** | 0.66 | 0.14 | 4.81 | 33.88 | <0.001 | 0.37 | 0.95 | 17.75 | 1 | <0.001 |
| **Trial number^1^** | 0.04 | 0.06 | 0.85 | 30.97 | 0.4 | -0.08 | 0.16 | 0.56 | 1 | 0.452 |
| **Condition²:outcomeIDENTITY** | -0.65 | 0.26 | -2.19 | 217.53 | 0.029 | -1.17 | -0.15 | 7.08 | 2 | 0.029 |
| **Condition²:outcomeLOCATION** | -0.1 | 0.26 | -0.11 | 213.88 | 0.909 | -0.59 | 0.41 |  |  |  |

Notes: ^1^Reference category: Eye-contact

^2^z-transformed

Table S2. Results of the LMM analyzing z-scored latency to the first fixation on the target object during the outcome phase. The table reports parameter estimates (Estimate ± SE), t values, approximate degrees of freedom (df), and p values from lmerTest, along with 95 % confidence intervals (CIs) and likelihood ratio test (LRT) results.

|  | **Estimate** | **SE** | **t** | **Df** | **p** | **LowerCI** | **UpperCI** | **Chi2** | **df_LRT** | **p_LRT** |
| --- | --- | --- | --- | --- | --- | --- | --- | --- | --- | --- |
| **(Intercept)** | 0.53 | 0.14 | 3.13 | 156.98 | 0.002 | 0.24 | 0.81 |  |  |  |
| **Conditition^2^** | -0.31 | 0.2 | -1.31 | 134.41 | 0.194 | -0.69 | 0.08 |  |  |  |
| **outcomeIDENTITY** | -0.46 | 0.18 | -2.2 | 194.16 | 0.029 | -0.8 | -0.11 |  |  |  |
| **outcomeLOCATION** | -0.25 | 0.19 | -1.11 | 90.42 | 0.271 | -0.63 | 0.12 |  |  |  |
| **object_pos_outcomeup** | -0.51 | 0.13 | -3.95 | 27.51 | <0.001 | -0.76 | -0.26 | 11.87 | 1 | 0.001 |
| **Trial number**^1^ | -0.11 | 0.06 | -1.76 | 208.1 | 0.08 | -0.24 | 0 | 3.51 | 1 | 0.061 |
| **Condition²:outcomeIDENTITY** | 0.77 | 0.26 | 2.49 | 179.44 | 0.014 | 0.25 | 1.26 | 8.87 | 2 | 0.012 |
| **Condition²:outcomeLOCATION** | 0.17 | 0.26 | 0.52 | 181.31 | 0.6 | -0.31 | 0.69 |  |  |  |

Notes: ^1^Reference category: Eye-contact

^2^z-transformed

Table S3. Results of the LMM analyzing z-scored sample-based looking time to the target object during the outcome phase. The table reports parameter estimates (Estimate ± SE), t values, approximate degrees of freedom (df), and p values from lmerTest, along with 95 % confidence intervals (CIs) and likelihood ratio test (LRT) results.

|  | **Estimate** | **SE** | **t** | **df** | **p** | **LowerCI** | **UpperCI** | **Chi2** | **df_LRT** | **p_LRT** |
| --- | --- | --- | --- | --- | --- | --- | --- | --- | --- | --- |
| **(Intercept)** | -0.57 | 0.14 | -3.64 | 153.42 | <0.001 | -0.84 | -0.29 |  |  |  |
| **Conditition^1^** | 0.23 | 0.18 | 1.11 | 223.59 | 0.269 | -0.1 | 0.58 |  |  |  |
| **outcomeIDENTITY** | 0.44 | 0.17 | 2.25 | 230.99 | 0.025 | 0.1 | 0.77 |  |  |  |
| **outcomeLOCATION** | 0.26 | 0.18 | 1.36 | 218.73 | 0.176 | -0.09 | 0.59 |  |  |  |
| **object_pos_outcomeup** | 0.64 | 0.13 | 4.63 | 34.065 | <0.001 | 0.38 | 0.87 | 17.63 | 1 | <0.001 |
| **Trial number^2^** | 0.01 | 0.06 | 0.2 | 32.26 | 0.842 | -0.11 | 0.12 | 0.01 | 1 | 0.934 |
| **Condition²:outcomeIDENTITY** | -0.65 | 0.25 | -2.31 | 230.38 | 0.022 | -1.12 | -0.16 | 6.64 | 2 | 0.036 |
| **Condition²:outcomeLOCATION** | -0.23 | 0.25 | -0.85 | 219.78 | 0.397 | -0.7 | 0.23 |  |  |  |

^1^Reference category: Eye-contact

^2^z-transformed

Table S4. Results of GAMM01 analyzing baseline-corrected pupil size during the action phase.

| *Parametric coefficients* |  |  |  |  |
| --- | --- | --- | --- | --- |
| Term | Estimate | SE | t | p |
| (Intercept) | -24.13 | 34.75 | -0.69 | 0.487 |
| Conditition^1^ | -53.88 | 48.96 | -1.1 | 0.271 |

| *Smooth terms* |  |  |  |  |
| --- | --- | --- | --- | --- |
| Smooth Term | edf | Ref.df | F | p |
| s(time):conditionCOM | 12.67 | 14.88 | 4.74 | <0.001 |
| s(time):conditionNCOM | 13.14 | 15.32 | 3.41 | <0.001 |
| s(Xgaze,Ygaze) | 23.7 | 27.12 | 97.94 | <0.001 |
| s(time,Event) | 3,323.07 | 3,502.00 | 155.8 | <0.001 |
| s(time,subject) | 32.56 | 314 | 0.12 | <0.001 |

Notes: Reference category: Eye-contact

Table S5. Demographic information of the tested dogs.

| Dog_ID | Breed | Sex | Neutered | Birth date | Age (in months) |
| --- | --- | --- | --- | --- | --- |
| subj.01 | Mix | m | yes | 01/06/2022 | 20 |
| subj.02 | Mix | m | yes | 13/11/2015 | 99 |
| subj.03 | Mix | m | yes | 28/04/2019 | 57 |
| subj.04 | Mix | f | no | 01/10/2018 | 64 |
| subj.05 | Mix | m | yes | 17/04/2020 | 45 |
| subj.06 | Mix | m | yes | 01/02/2019 | 60 |
| subj.07 | Mix | m | yes | 09/07/2015 | 103 |
| subj.08 | Collie | m | no | 12/04/2019 | 58 |
| subj.09 | Mix | f | no | 20/02/2022 | 23 |
| subj.10 | Leisha Dog | f | no | 20/10/2019 | 51 |
| subj.11 | Labrador Retriever | f | yes | 14/08/2018 | 66 |
| subj.12 | Border Collie | m | yes | 15/03/2019 | 59 |
| subj.13 | Border Collie | f | yes | 17/02/2017 | 84 |
| subj.14 | Australian Shepherd | m | yes | 26/10/2016 | 88 |
| subj.15 | Mix | f | yes | 06/11/2020 | 39 |
| subj.16 | Flat Coated Retriever | f | no | 06/02/2016 | 97 |
| subj.17 | Welsh Springer Spaniel | f | no | 17/06/2022 | 20 |
| subj.18 | Labrador Retriever | f | no | 18/08/2020 | 42 |
| subj.19 | Mix | f | yes | 07/09/2017 | 78 |
| subj.20 | Magyar Vizsla | f | no | 20/05/2022 | 22 |
| subj.21 | Nova Scotia Duck Tolling Retriever | f | no | 03/03/2018 | 72 |
| subj.22 | Fox Terrier | m | yes | 15/01/2021 | 38 |
| subj.23 | Mix | m | yes | 01/09/2018 | 67 |
| subj.24 | Mix | m | yes | 01/06/2019 | 58 |
| subj.25 | Australian Shepherd | m | no | 31/10/2021 | 29 |
| subj.26 | Magyar Vizsla | m | no | 20/02/2021 | 37 |
| subj.27 | Mix | f | no | 01/05/2020 | 47 |
| subj.28 | Mix | m | yes | 02/06/2021 | 34 |
| subj.29 | Mix | f | no | 10/01/2019 | 64 |
| subj.30 | Border Collie | f | no | 05/10/2018 | 67 |
| subj.31 | Border Collie | m | no | 02/06/2023 | 11 |
| subj.32 | Australian Shepherd | m | no | 18/08/2016 | 94 |
| subj.33 | Australian Shepherd | f | no | 13/03/2022 | 28 |
| subj.34 | Labrador Retriever | f | no | 05/05/2020 | 50 |
| subj.35 | Mix | f | no | 01/04/2022 | 28 |
| subj.36 | Australian Shepherd | m | no | 20/04/2019 | 64 |

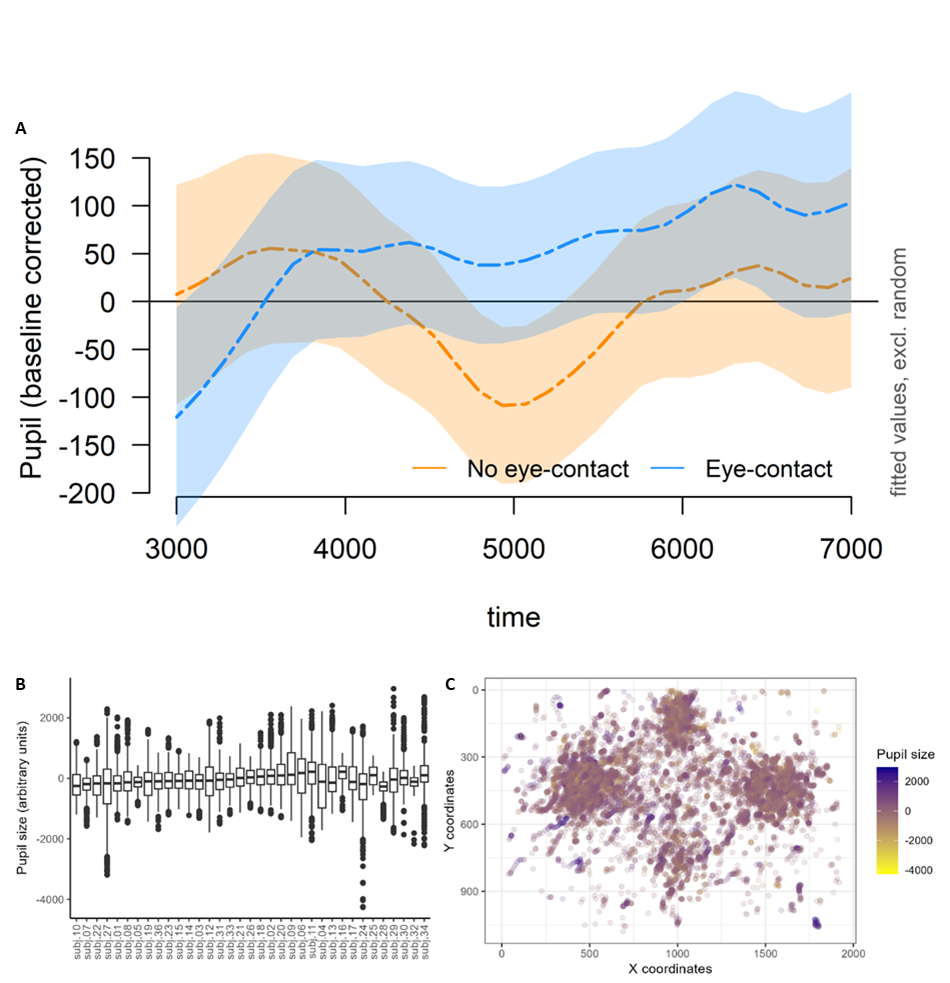
Figure S1. (A) Summed GAMM effects of pupil size as a function of time during the action phase. Model-predicted, baseline-corrected pupil size trajectories are shown for the eye-contact and no-eye-contact conditions, with shaded areas indicating approximate standard errors. (B) Distribution of baseline-corrected pupil size values across subjects (after removal of blink artefacts). (C) Preprocessed and baseline-corrected pupil size values plotted against gaze position (X and Y screen coordinates).

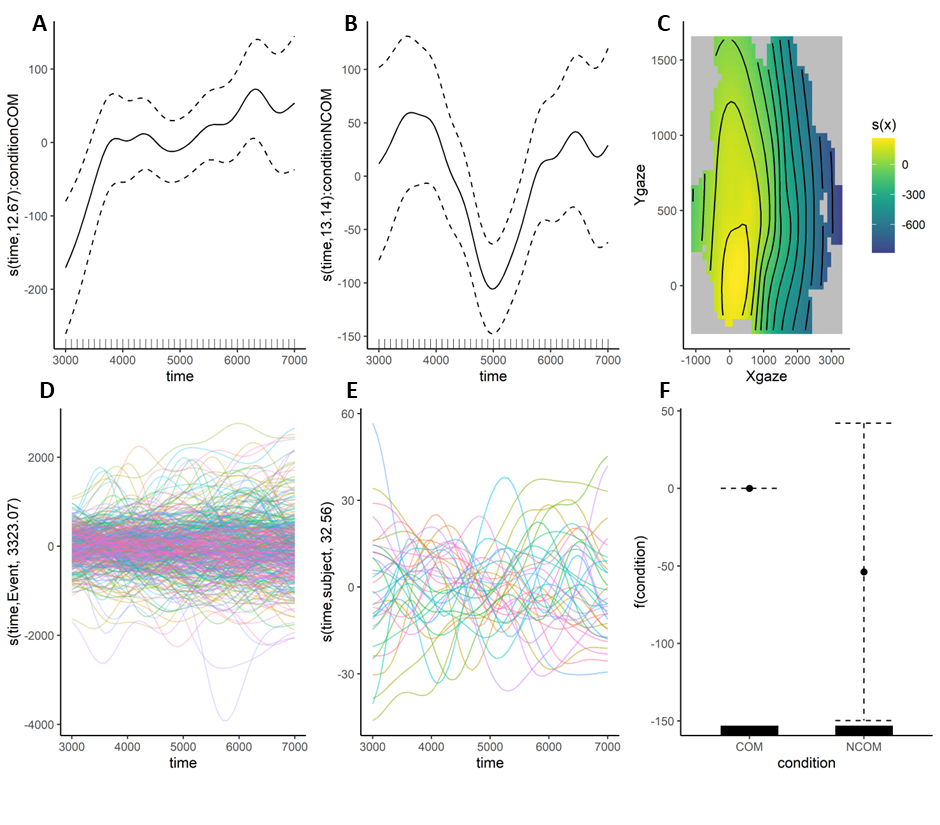

**Figure S2.** Partial and summed effects of GAMM01 for the action phase. (A–B) Smooth terms for time in the eye-contact (COM) and no-eye-contact (NCOM) conditions. (C) Two-dimensional smooth of gaze position (X and Y coordinates). (D) Random smooths for event-level effects. (E) Random smooths for individual subjects. (F) Parametric effect of condition, showing estimated differences between COM and NCOM.

**-**

**Video S1.** The video shows a complete testing trial from the eye-contact condition. The actor shifts her gaze between the dog and the object during the action phase, followed by the occlusion and outcome phases. A moving dot is superimposed to indicate the dog’s gaze position throughout the trial.

**Video S2.** This video displays a complete testing trial from the no-eye-contact condition. The actor avoids direct gaze and alternates her attention between the object and her original side-facing position. A moving dot is superimposed to indicate the dog's gaze position throughout the trial.
